# Fitness effects of mutations to SARS-CoV-2 proteins

**DOI:** 10.1101/2023.01.30.526314

**Authors:** Jesse D. Bloom, Richard A. Neher

## Abstract

Knowledge of the fitness effects of mutations to SARS-CoV-2 can inform assessment of new variants, design of therapeutics resistant to escape, and understanding of the functions of viral proteins. However, experimentally measuring effects of mutations is challenging: we lack tractable lab assays for many SARS-CoV-2 proteins, and comprehensive deep mutational scanning has been applied to only two SARS-CoV-2 proteins. Here we develop an approach that leverages millions of publicly available SARS-CoV-2 sequences to estimate effects of mutations. We first calculate how many independent occurrences of each mutation are expected to be observed along the SARS-CoV-2 phylogeny in the absence of selection. We then compare these expected observations to the actual observations to estimate the effect of each mutation. These estimates correlate well with deep mutational scanning measurements. For most genes, synonymous mutations are nearly neutral, stop-codon mutations are deleterious, and amino-acid mutations have a range of effects. However, some viral accessory proteins are under little to no selection. We provide interactive visualizations of effects of mutations to all SARS-CoV-2 proteins (https://jbloomlab.github.io/SARS2-mut-fitness/). The framework we describe is applicable to any virus for which the number of available sequences is sufficiently large that many independent occurrences of each neutral mutation are observed.

---

The rapid evolution of SARS-CoV-2 has led to the emergence of viral variants with enhanced transmissibility, escape from therapeutics, or reduced recognition by immunity [1, 2]. To anticipate and mitigate this evolution, the scientific community has launched efforts to assess the risk of new viral variants [3] and create therapeutics that target constrained regions of the virus where resistance is less likely to evolve [4, 5, 6]. Both efforts require determining how specific mutations affect viral fitness. Unfortunately, experimentally measuring the effects of mutations is challenging for most SARS-CoV-2 proteins. For spike, tractable lab assays have identified key functional and antigenic mutations [1, 7], and enabled deep mutational scanning measurements of how most mutations affect receptor binding, cellular infection, and antibody recognition [8, 9, 10, 11]. These experimental data are valuable for assessing new spike variants [3, 12, 13] and designing antibody therapeutics with greater resistance to escape [14, 15, 16]. But most SARS-CoV-2 proteins lack tractable lab assays, despite contributing to viral fitness [17, 18, 19] and being targets of efforts to develop anti-viral drugs [20]. The only non-spike SARS-CoV-2 protein with large-scale experimental measurements of mutation effects is Mpro [21, 22].

An alternative to experiments is to estimate effects of mutations by analyzing natural viral sequences. The amount of data available for such analyses has increased dramatically over the last few years with the sequencing of SARS-CoV-2 from millions of human infections. So far analyses of these sequences have focused on analyzing expanding viral clades to identify mutations that mediate immune escape or increase transmissibility [23, 24, 25]. The basic idea is that mutations that repeatedly appear near the base of clades that increase in relative frequency are likely beneficial to the virus. However, only a small minority of all possible mutations are beneficial, with most being nearly neutral or deleterious. For purposes such as identifying constrained drug targets or understanding the function of viral proteins, it is important to estimate the effects of neutral or deleterious mutations as well as beneficial ones. Other studies have analyzed broader alignments of coronaviruses substantially diverged from SARS-CoV-2 [26, 27], but the resulting estimates are limited by sparse sampling and possible changes in the impacts of some mutations across divergent viruses.

Here we develop a new approach that uses natural sequences to estimate the effects of mutations. Our basic insight is that there are now so many SARS-CoV-2 sequences that all non-deleterious single-nucleotide mutations are expected to independently occur many times along the observed phylogenetic tree. We therefore first calculate the number of expected observations of independent occurrences of each mutation based on the neutral mutation rate of SARS-CoV-2. We then compare these expected observations to the actual observations in the SARS-CoV-2 tree to estimate the effect of each mutation. The resulting estimates correlate well with existing deep mutational scanning data. Most viral proteins have regions under strong selective constraints. However some accessory proteins show only weak selection against amino-acid and even stop-codon mutations. Overall, our work demonstrates a new approach to determine the effects of mutations, and provides detailed maps of mutational effects across the SARS-CoV-2 proteome.

## Results

### Mutation effects from actual versus expected counts

To determine how many times each mutation is expected to be observed, we used the pre-built UShER tree [28, 29, 30] of *∽*7-million public SARS-CoV-2 sequences to count nucleotide mutations at four-fold degenerate sites [Figure 1A; 31]. Because mutations at such sites never alter the amino-acid sequence, these counts reflect the mutation process in the absence of protein-level selection (see below for caveats about nucleotide-level selection). The expected counts of a mutation from nucleotide *x* to *y* is simply the average count of this type of mutation across all four-fold degenerate sites with parental identity *x*. Importantly, we count independent *occurrences* of each mutation along the branches of the tree, not the sequences with the mutation in the final alignment (Figure 1A. We also compute expected counts separately for each SARS-CoV-2 clade to account for shifts in mutation spectrum [31, 32], and apply quality-control steps to remove spurious mutations (see Methods).

**Figure 1.**
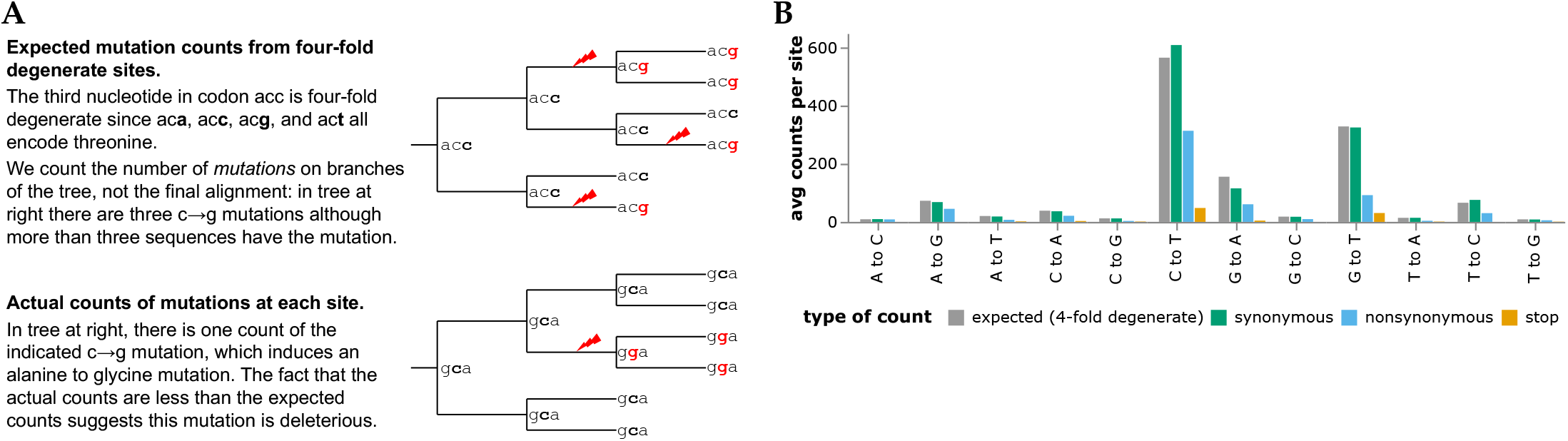
Expected versus actual counts of mutations. **(A)** The number of expected counts of each type of nucleotide mutation is computed from four-fold degenerate sites, and then compared the actual counts of each mutation. **(B)** Expected versus actual counts for each nucleotide mutation type aggregated across all viral clades and averaged across all sites where the mutation is four-fold degenerate, synonymous (including four-fold degenerate), nonsynonymous, or introduces a stop codon. See https://jbloomlab.github.io/SARS2-mut-fitness/avg_counts.html for an interactive version of panel B that enables mouseovers to read off specific values.

The expected counts per mutation vary with mutation type, ranging from *∽*565 for C*→*T to only *∽*9 for T*→*G mutations (Figure 1B). This variation is because the SARS-CoV-2 mutation spectrum is highly biased towards specific mutation types [31, 32, 33, 34]. However, because there are so many SARS-CoV-2 sequences we are able to estimate the rate of even the rarest mutation types with high accuracy [31]. For instance, there are *∽*1.9 *×* 10^4^ observed occurrences of T*→*G mutations across all *∽*2,100 four-fold degenerate sites with a parental identity of T, which is enough to estimate the T*→*G mutation rate (and therefore the expected counts of each mutation) with high accuracy.

We compared the expected counts to the actual observed counts of mutations averaged across sites (Figure 1). For synonymous mutations, the expected and actual counts are similar. But for nonsynonymous and especially stop-codon mutations, the actual counts are substantially lower than the expected counts, reflecting purifying selection for protein function.

The ratio of actual to expected counts for each mutation is related to its effect on viral fitness. The intuition is straightforward: mutations arise at all sites, but viruses with deleterious mutations are less likely to transmit and be observed in sequencing of human SARS-CoV-2. Therefore, the ratio of actual to expected counts will be one for neutral mutations, and less than one for deleterious mutations. In the Methods and Appendix, we show that under plausible assumptions about SARS-CoV-2 evolution and sampling intensity (fraction of viruses sequenced), the fitness cost of a deleterious mutation scales roughly inversely with the ratio of actual to expected counts for mutations with costs greater than a few percent. A key result is the dependence on sampling intensity: if all human SARS-CoV-2 were sequenced even deleterious mutations would have a high chance of being sampled and we would need to study the subsequent spread of the mutations to assess their fitness. But the actual sampling intensity is *∽*0.1%, since there are *∽*7-million publicly available SARS-CoV-2 sequences and the total number of human infections is now probably roughly on par with the total global population of *∽*8-billion people. At this sampling intensity, the number of times a mutation is observed reflects more subtle reductions in transmission efficiency. We quantify the effect of each mutation as the logarithm of the ratio of actual to expected counts after summing counts for all nucleotides that encode the relevant amino-acid. The statistical noise is greater for mutations with fewer expected counts: the figures in this paper show mutations with *≥* 10 expected counts unless otherwise noted, with legends linking to interactive plots that enable adjustment of this threshold.

### Mutation-effect estimates are robust to subsampling, with some evidence of epistasis in spike

We computed the correlations among mutation-effect estimates made using subsets of SARS-CoV-2 sequences from different viral clades or geographic locations. These estimates were well correlated, with some modest variation in estimates across sequence subsets (Figure 2A,B).

**Figure 2.**
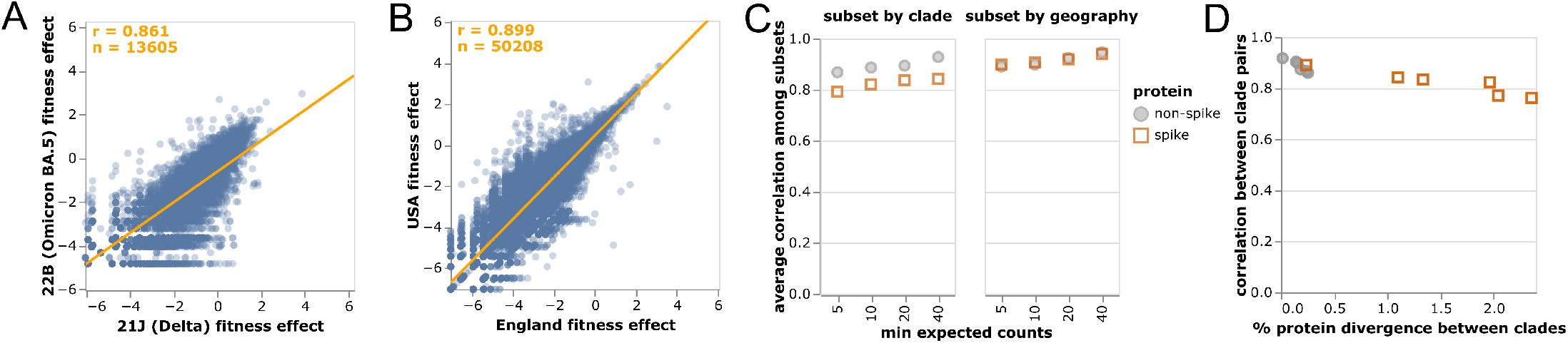
Correlations of mutation fitness effect estimates made using subsets of natural sequences. Correlations between estimates made **(A)** just using sequences from the Delta or Omicron BA.5 clades or **(B)** just from the USA or England. Each point is an amino-acid mutation, the orange line is a least-squares regression, and orange text at upper left shows the number of mutations and Pearson correlation coefficient. Only mutations with at least 10 expected counts are shown, which is why panels have different numbers of mutations shown (sequence subsets vary in size). Different subset size are also the reason why the regression line in (A) deviates from the identity *x* = *y*. **(C)** Correlations between clade or geography subsets become higher with an increasingly large threshold for minimum expected counts. Spike mutations have a worse correlation when subsetting by viral clade (plot shows average correlation over all pairwise combinations of Delta, BA.1, BA.2, and BA.5), but not when subsetting by geography (USA or England). **(D)** Correlations in estimated mutation effects decline for clades with higher protein divergence, with the effect most noticeable for spike since spike is more diverged among SARS-CoV-2 clades than other viral proteins. See https://jbloomlab.github.io/SARS2-mut-fitness/clade_corr_chart.html and https://jbloomlab.github.io/SARS2-mut-fitness/subset_corr_chart.html for versions of A and B that include all viral clades with at least 500,000 total expected counts (summed across all mutations) and have other interactive options.

The modest variation in estimates from different sequence subsets could have two causes: statistical noise due to finite mutation counts, or real shifts in mutation effects during SARS-CoV-2 evolution [35, 36]. To test for statistical noise, we computed correlations with different thresholds for how many expected counts are required before making an estimate for a mutation (Figure 2C). Correlations increased with this count threshold, consistent with reduced statistical noise for larger mutation counts. But the correlation for spike mutations was consistently lower for cross-clade but not cross-geography comparisons (Figure 2C). The lower cross-clade correlation for spike appears due to epistatic shifts in mutation effects [35, 36, 37, 38, 39] or changes in the selective landscape [40] between SARS-CoV-2 clades, since the correlation is lower between clades with higher spike divergence (Figure 2D). In particular, the interactive version of Figure 2A shows that mutations that are more beneficial in Omicron BA.5 than Delta are often antibody-escape mutations (eg, K444N or G446S in spike [12])—a result that makes sense, since newer variants like Omicron BA.5 are evolving under increased immune selection compared to earlier variants like Delta that circulated in a more immunologically naive population.

Despite evidence for some shifts in mutation effects in spike, for the rest of this paper we aggregate counts across viral clades to make a single estimate for each amino-acid mutation. The reason is that the accuracy of the estimates increases with the number of counts (Figure 2C), and several mutation types only have enough counts for reasonable estimates when aggregating across clades (Figure 1B). For the purposes of this paper, we deemed it preferable to have more accurate and comprehensive pan-SARS-CoV-2 estimates than noisier clade-specific estimates for fewer mutations. However, the interactive version of Figure 2A linked in the legend enables exploration of mutations with disparate estimates among clades.

An important question is whether the mutation fitness estimates are affected by noise from limited statistical sampling of mutations or whether sequencing errors and bioinformatic artefacts distort the estimates. To assess if this is the case, we repeated the entire fitness estimation using an even larger pre-built UShER mutation-annotated tree of all *∽*14-million SARS-CoV-2 sequences in GISAID [41] as of March-29-2023. There is an extremely high correlation between fitness estimates made using the *∽*7-million publicly available sequences and the larger GISAID tree (Figure S1). This concordance indicates that the set of *∽*7-million public sequences is large enough that doubling the data does not appreciably shift the estimates, and so throughout this paper we use that sequence set due to our preference for publicly available data. Furthermore, fitness estimates using sequences from specific countries (USA and England, Figure 2B) are also highly concordant, suggesting that sequencing and bioinformatic workflows are not driving the signal. Lastly, positions known to be under strong constraint *a priori* (e.g. start codons, the ribosomal slippage site) typically have no or only few mutations, suggesting that sequencing errors in consensus sequences are rare.

### Structural and non-structural proteins are under strong purifying selection, but most accessory proteins are not

The distributions of mutation effects concur with biological intuition about how different classes of mutations impact protein function. Most synonymous mutations are nearly neutral, most stop codons are highly deleterious, and amino-acid mutations range from slightly beneficial to highly deleterious (Figure 3A). The handful of synonymous mutations with highly deleterious effects are in either regions of known non-coding constraint (e.g., the ORF1ab ribosomal slippage site [43]) or two regions in the center of E and the end of M (Figure S2).

**Figure 3.**
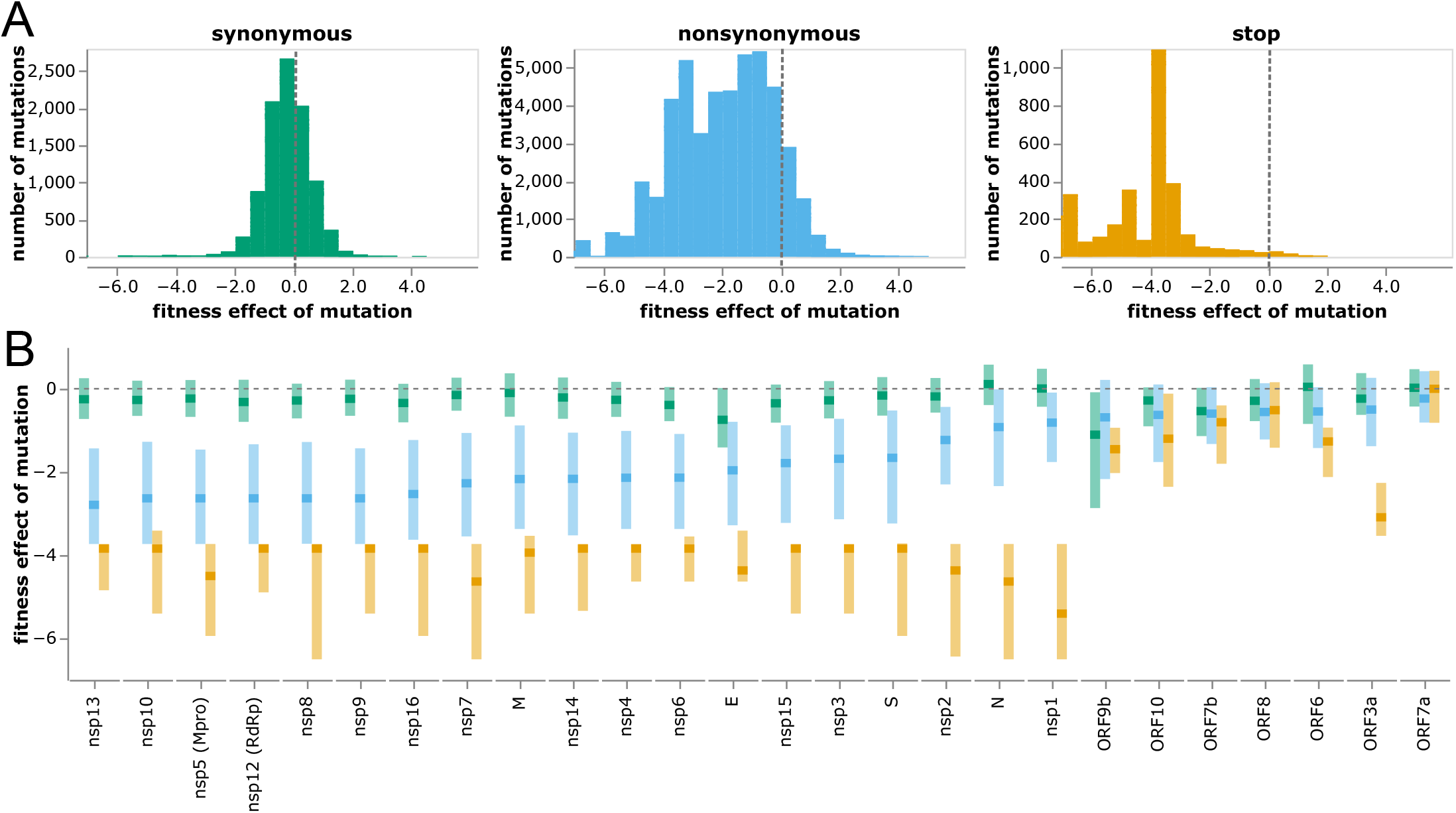
Distribution of effects of different classes of mutations. **(A)** Histograms of effects of synonymous, nonsynonymous, and stop-codon mutations across all viral genes. Neutral mutations have effects of zero (dashed gray vertical lines), and deleterious mutations have negative effects. **(B)** Effects of each class of mutation for each viral gene. Dark squares indicate the median effect, and the lighter rectangles span the interquartile range. Mutation types are color-coded as in panel A. The apparent constraint on synonymous mutation in ORF9b is probably because this gene is encoded in an overlapping reading frame with N [42]. See https://jbloomlab.github.io/SARS2-mut-fitness/effects_histogram.html and https://jbloomlab.github.io/SARS2-mut-fitness/effects_dist.html for plots that allow adjustment of the expected-count cutoff and other interactive options (such as separate histograms for each gene). See Figure S3 for a version of panel B with genes ordered by genomic position rather than constraint on nonsynonymous mutations.

To investigate differences in functional constraint among viral proteins, we computed the distributions of mutation effects separately for each gene (Figure 3B and S3). SARS-CoV-2 proteins are grouped into three categories: nonstructural (or nsp) proteins, structural proteins (spike, M, N, and E), and accessory proteins (names prefixed with “ORF”) [44]. The nonstructural and structural proteins are essential, and these proteins show strong selection against stop codons and clear although variable purifying selection against amino-acid mutations (Figure 3B and S3; e.g., nsp13 is under stronger protein-level constraint than nsp1).

However, most accessory proteins are under little constraint (Figure 3B and S3). Stop-codon and amino-acid mutations to ORF7a and ORF8 are not more deleterious than synonymous mutations (although recall that our estimates are only sensitive to fitness costs greater than a few percent). The lack of deleterious mutations to ORF8 is consistent with the fact that viruses with deletions in this gene have spread in humans [45] and that major variants had stop codons early in ORF8. Indeed, the loss of accessory proteins such as ORF8 appears to occur with some regularity during the early evolution of non-human viruses in humans [46]. The only accessory protein under strong purifying selection against stop codons is ORF3a (Figure 3B), for which stop codons in the first 240 residues are clearly deleterious (Figure S4). These observations concur with experiments showing SARS-CoV-2 is attenuated by deletion of ORF3a but there is little effect of deleting ORF6, ORF7a, or ORF8 [19, 47, 48]. However, ORF3a’s function must be relatively insensitive to its protein sequence, since other than selection against stop codons there is only amino-acid level constraint at a few sites like 135 and 138 (Figure S4). Observations such as these could help guide experimental studies to better understand protein function.

### Mutation-effect estimates correlate with experiments

We examined how the mutation effects estimated using our approach compare with prior high-throughput deep mutational scanning measurements. For spike, two distinct experimental methodologies have been used to characterize large numbers of mutations: yeast display of the receptor-binding domain (RBD) [8, 49] and spike pseudotyped lentiviruses [9]. For Mpro (also known as nsp5 or 3CLpro), two different labs have performed deep mutational scanning using the same basic methodology of assaying protease cleavage in yeast [21, 50, 22].

For spike, our estimates from natural sequences correlate with the experiments almost as well as the two experimental methodologies correlate with each other (Figure 4A), with Pearson correlations of 0.66 between the estimates and experiments versus 0.72 between the two experiments. Neither experiment fully captures how mutations affect viral fitness, since both RBD yeast display and lentiviral pseudotyping are imperfect proxies for spike function during actual human infections. Therefore, it is unclear how much the differences between the mutation-effect estimates and experiments are due to noise in the estimates versus limitations of the experiments. However, the fact that the estimates correlate with the experiments almost as well as the experiments correlate with each (Figure 4A) suggests the estimates are of comparable quality to experimental measurements. At least some of the mutations with the greatest divergence between our estimates and the deep mutational scanning likely represent experimental artifacts. For instance, P527L, which is favorable in the RBD deep mutational scan but deleterious in the sequence-based estimates and full-spike scan, is at the C-terminus of the yeast-displayed RBD [8] where it may adopt a non-native conformation.

**Figure 4.**
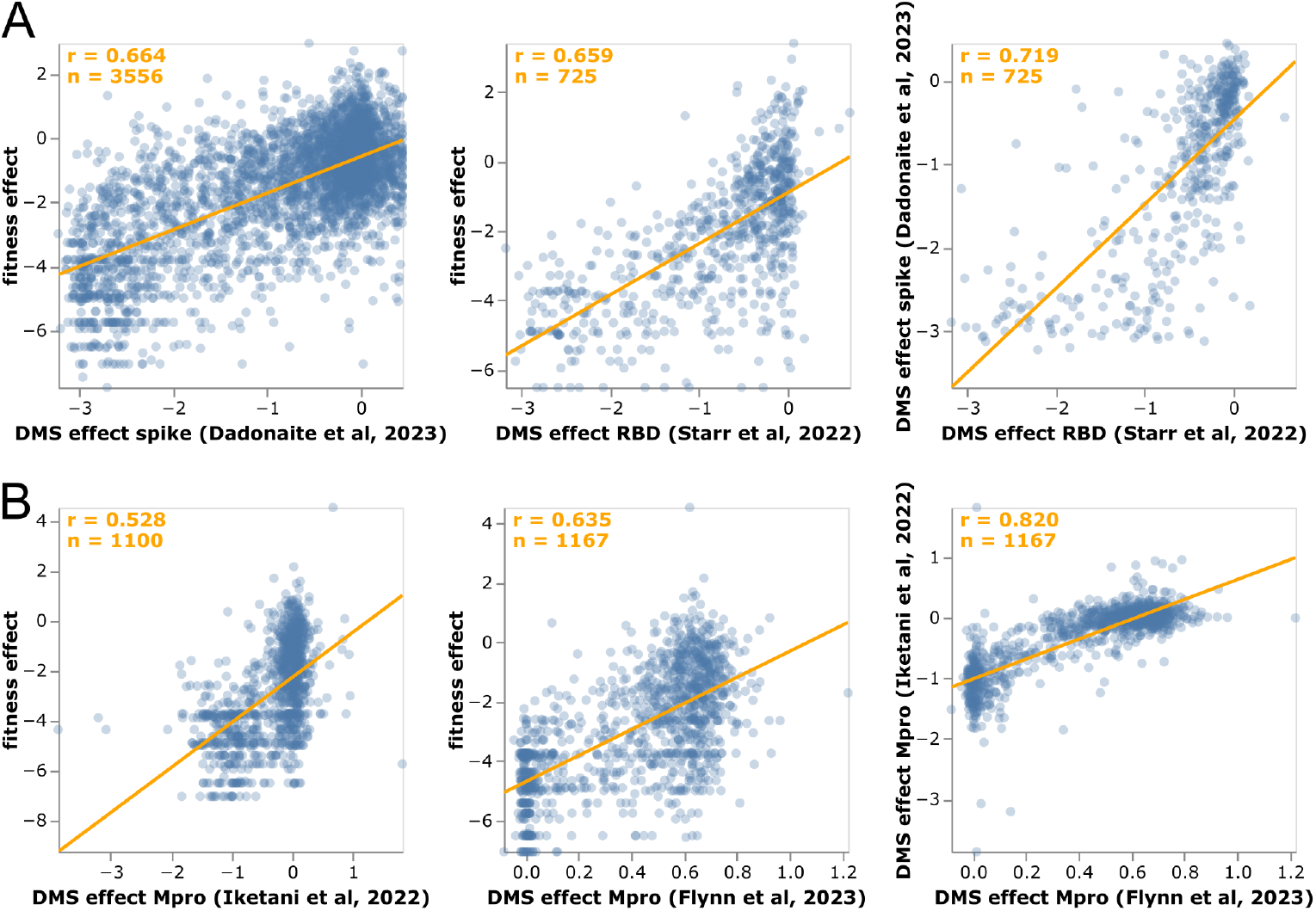
Correlation of mutation-effect estimates with experimental deep mutational scanning measurements for **(A)** the full spike [9] or its RBD [49], and **(B)** Mpro [50, 22]. Each point is an amino-acid mutation, the orange line is a least-squares regression, and orange text in the upper left shows the number of mutations and Pearson correlation coefficient. Each sub-panel shows a different set of mutations (depending on which mutations were measured in that experiment). See https://jbloomlab.github.io/SARS2-mut-fitness/dms_S_corr.html and https://jbloomlab.github.io/SARS2-mut-fitness/dms_nsp5_corr.html for plots that also show the Mpro dataset from [21] and have various interactive options. The plots in this figure show the average of the multiple phenotypes measured in the deep mutational scanning of [49]; see https://jbloomlab.github.io/SARS2-mut-fitness/dms_S_all_corr.html for each phenotype separately. This figure only shows mutations with at least 20 expected counts, which is higher than the threshold of 10 used in most of the rest of this paper (this threshold can be adjusted in the interactive plots).

The sequence-based estimates for Mpro also correlate with the deep mutational scans for that protein, although in this case the experiments correlate substantially better with each other than with our estimates (Figure 4B). However, the Mpro experiments all use a similar yeast-based methodology [21, 50, 22] that fails to capture significant aspects of Mpro’s function during human infections. For instance, a stop codon at Q306 is well tolerated in the deep mutational scans but extremely disfavorable in our sequence-based estimates, and such a mutation would clearly be highly deleterious to actual virus as it would truncate the polyprotein. Similarly, K61N is well tolerated in the deep mutational scans but extremely disfavorable in our estimates, probably because in the full viral polyprotein this residue mediates important interactions between Mpro and nsp7-10 [51]

### Mutation-effect estimates better capture functional constraint than dN/dS ratios or predictions from other methods

A longstanding approach for analyzing protein constraint is to compare rates of nonsynonymous (dN) and synonymous (dS) substitutions at each site [52, 53]. These dN/dS ratios can be calculated by counting mutations or using phylogenetic substitution models [53, 54]. A limitation of dN/dS ratios is they cannot be interpreted in terms of the fitness effects, since they simply represent the relative rate of amino-acid substitution rather than the effects of specific mutations [55, 56]. Nonetheless, we can compare dN/dS ratios to our mutation-effect estimates as measures of the average constraint at each site. The mutation-effect estimates greatly outperform dN/dS ratios as a measure of site-level constraint as assessed by correlation with deep mutational scanning experiments (Figure S5). The reason is in part because some aspects of functional constraint cannot be captured by a dN/dS ratio. For instance, the ACE2-affinity enhancing spike mutation N501Y arose in several SARS-CoV-2 variants early in the pandemic, and has since remained fixed due its importance for receptor binding [49]. Our mutation-effect estimates correctly reflect that site 501 is strongly prefers tyrosine, but the site has a high dN/dS ratio due to the early convergent evolution of this site to that preferred amino acid.

Our mutation-effect estimates also correlate better with deep mutational scanning experiments than predictions from two algorithms trained to learn epistatic models of mutation effects from phylogenetically broader but more sparsely sampled sequence data [27, 26] (Figure S6). Our estimates also correlate better with experiments than predictions by a machine-learning algorithm that integrates sequence and epidemiological data [25] (Figure S6). These results suggest that our straightforward approach of directly reading out the effects of mutations from their actual versus expected counts can outperform much more complex models when millions of sequences are available.

### Fixed mutations tend to have beneficial or neutral effects

Amino-acid mutations that have fixed in at least one viral clade are estimated to mostly have neutral or beneficial effects, whereas most other mutations are deleterious (Figure S7). This fact is unsurprising: viral lineages that expand into new clades do so because they have acquired beneficial mutations while avoiding deleterious ones [57, 58, 59]. But the fact that the beneficial effects of fixed mutations are correctly estimated by our approach, which simply counts mutation occurrences and does not incorporate information on lineage size, demonstrates such mutations occur independently in many viral lineages that are more successful than average.

Most fixed mutations are estimated to be beneficial regardless of whether estimates are made using all viral clades, or just clades that did not fix the mutation (Figure S8). However, a few beneficial fixed mutations show epistatic entrenchment [38, 60] in the sense that they are not particularly beneficial in clades in which they did not fix (Figure S8). The most striking example is S373P in spike, which has experimentally been shown to be neutral or slightly deleterious in pre-Omicron clades, but strongly beneficial in the Omicron clades in which it fixed [49, 36].

### Interactive exploration of amino-acid fitnesses

To enable easy access to the mutation-effect estimates, we created interactive plots to enable exploration of the data for each protein. A static view of one of these plots is in Figure 5; see https://jbloomlab.github.io/SARS2-mut-fitness for interactive versions for all proteins. These plots enable both high-level inspection of functional constraint across each protein, and detailed interrogation of the effects of specific mutations.

**Figure 5.**
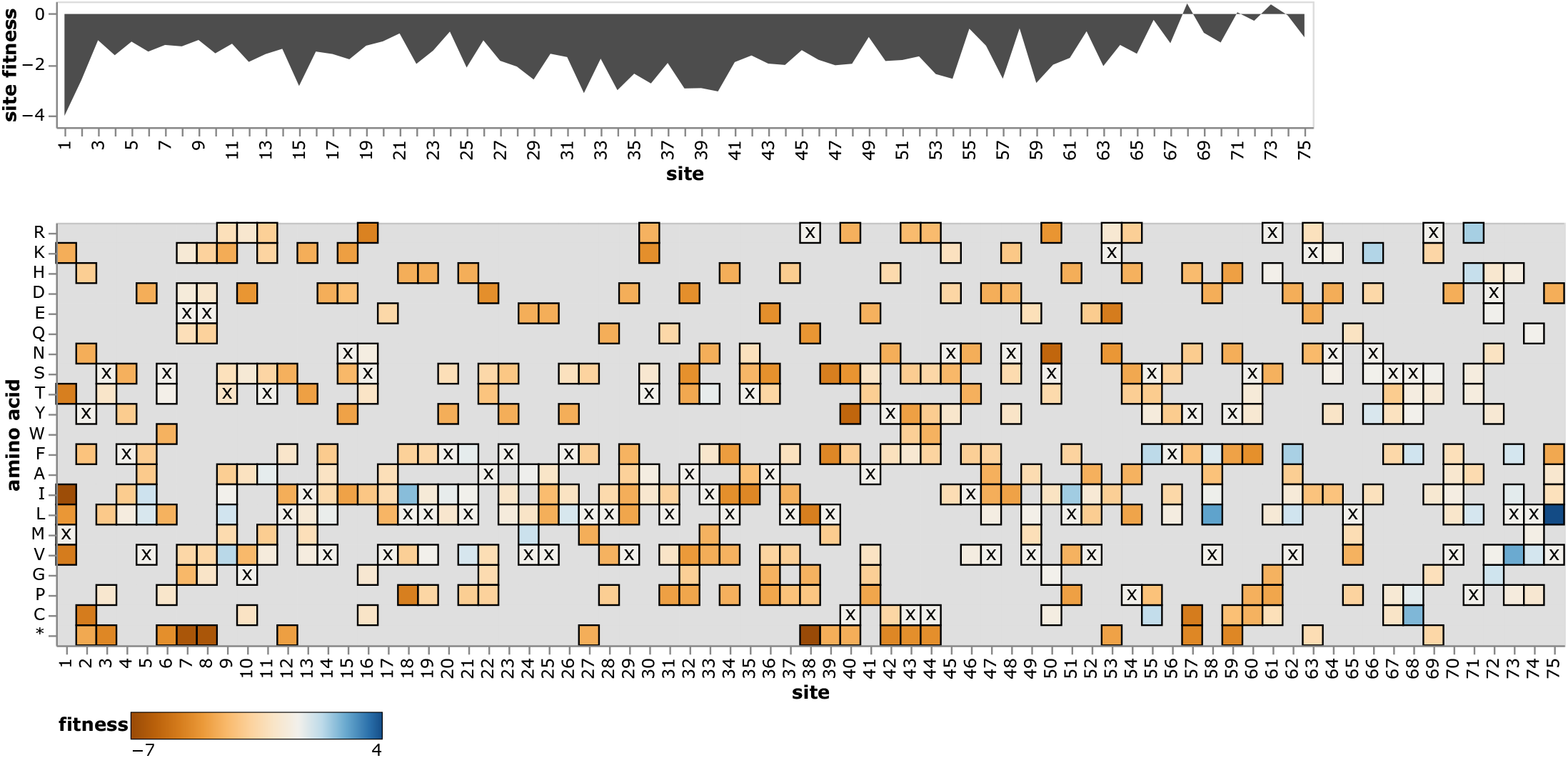
Effects of amino-acid mutations to E protein. The area plot at top shows the average effects of mutations at each site, and the heatmap shows the effects of specific amino acids, with **x** denoting the amino-acid identity in the Wuhan-Hu-1 strain. See https://jbloomlab.github.io/SARS2-mut-fitness/E.html for an interactive version of this plot that enables zooming, mouseovers, adjustment of the minimum expected count threshold, and layering of stop codon effects on the site plot. See https://jbloomlab.github.io/SARS2-mut-fitness for comparable interactive plots for all SARS-CoV-2 proteins.

## Discussion

Enough SARS-CoV-2 viruses have now been sequenced that many independent occurrences of every tolerated single-nucleotide mutation have been observed along the viral phylogeny. Here we have described a new approach that leverages this fact to estimate the effects of these mutations. In essence, we treat natural evolution as a deep mutational scan, with the millions of publicly available SARS-CoV-2 sequences providing a readout of this experiment. The key is simply to calculate how many times each mutation has been “tested” along the history of sampled viral sequences, and compare that expectation to the actual observations of the mutation among viruses sufficiently fit to have been sequenced in actual human infections.

The resulting estimates of mutational effects are robust to subsetting on specific viral clades or geographies, and correlate well with experimental measurements. In broad strokes, the mutation effects illuminate patterns of constraint: for instance, there is strong selection on structural and non-structural proteins, but only limited purifying selection on the accessory proteins.

However, the real value of our approach is in the detailed maps of effects of specific mutations to all viral proteins, including proteins with poorly understood functions not easily characterized in the lab. These maps will be of value for designing drugs that target constrained sites, interpreting the consequences of mutations observed during viral surveillance, and guiding experiments to mechanistically characterize protein function.

There are several caveats to our approach. First, because the number of observations of any given mutation is small compared to the millions of SARS-CoV-2 sequences being analyzed, our approach requires careful quality control to remove sequencing errors. Second, we assume the rate of each type of nucleotide mutation is uniform across the viral genome, and neglect higher-order context that may influence mutation rate [61, 62]. Likewise, we neglect constraint on nucleotide identity beyond the encoded protein sequence [63, 64]—although this probably has only a minor effect, since our analyses show just a handful of synonymous sites are under strong selection (Figure S2). Third, the exact relationship between the statistics we calculate and viral fitness depend on the fraction of all infections that are sequenced (sampling intensity) and viral population dynamics. Although we derive this relationship, we do not adjust for sampling intensity and population dynamics when estimating mutation effects. Fourth, we make a single estimate for each mutation across all SARS-CoV-2, neglecting the epistasis that can affect some mutations [35, 36]. Finally there are a few technical caveats to how we count mutations that are discussed in the Methods section.

Conceptually, our approach differs from prior methods that aim to identify beneficial SARS-CoV-2 mutations associated with viral clades that increase in frequency [23, 24, 25]. Those methods draw information primarily from what happens down-stream of a mutation. In contrast, we treat all mutations equivalently regardless of whether they are on a tip node or at the base of a large clade. Our approach is better for estimating effects of deleterious or nearly neutral mutations, but clade-growth methods may be better for beneficial mutations. In particular, clade size carries information beyond that contained in mutation counts alone (Figure S9). Hopefully future work can combine mutation-counting and clade-growth methods for even better estimates of SARS-CoV-2 mutation effects. Note our approach is conceptually similar to estimating fitness costs of HIV or polio mutations from mutation-selection balance in deep sequencing of intra-population viral quasispecies [65, 66], except we analyze mutation occurrences rather than frequencies to account for the phylogenetic structure and genetic hitchhiking that characterize global SARS-CoV-2 evolution.

The power of the approach we have described will increase with more viral sequencing. SARS-CoV-2 is the first virus with enough sequences that every tolerated mutation is observed multiple independent times. As costs drop, it is easy to imagine a future with even more viral sequences. As this occurs, viral genomic sequencing—which has traditionally been used primarily to track evolution and spread—will also become an increasingly precise tool to determine the effects of specific mutations.

## Methods

### Code and data availability

See the GitHub repository at https://github.com/jbloomlab/SARS2-mut-fitness for the computer code and processed data (eg, fitness estimates and mutation counts). That repository contains a README with links to specific data files as well as a description of the computational pipeline. See https://github.com/jbloomlab/SARS2-mut-fitness/blob/main/results/aa_fitness/aa_fitness.csv final estimates of amino-acid fitnesses across all clades; other intermediate data files are also provided in the GitHub repository. The specific version of the repository used for this paper is tagged as “bioRxiv-v2” on GitHub (https://github.com/jbloomlab/SARS2-mut-fitness/tree/bioRxiv-v2) The pipeline is fully reproducible, and is run using snakemake [67] with interactive plots rendered using altair [68].

The interactive plots are rendered at https://jbloomlab.github.io/SARS2-mut-fitness via GitHub pages.

### Versioning of analyses of different sequence sets

The figures in this manuscript show analyses of the set of all publicly available sequences as of May-11-2023. However, the pipeline can be run on different sequence sets. The sequence sets on which the analysis is currently run include all sets listed under the “mat_trees” key in https://github.com/jbloomlab/SARS2-mut-fitness/blob/main/config.yaml; these include the sets of all public sequences from several earlier dates (such as those available for the first version of this analysis), as well as the set of all sequences in GISAID as of March-29-2023. A version of the results for each sequence set is provided in the GitHub repository (https://github.com/jbloomlab/SARS2-mut-fitness) in subdirectories with names like “results_public_2023-05-11”, and the index page for the interactive plots (https://jbloomlab.github.io/SARS2-mut-fitness) links at the bottom to plots for each sequence set. The “current_mat” key in https://github.com/jbloomlab/SARS2-mut-fitness/blob/main/config.yaml specifies which sequence set is used to generate the main results that in the “results” subdirectory in the GitHub repository and are shown by default in the interactive plats; we anticipate periodically updating this to newer sequence sets as more sequences become available. See https://jbloomlab.github.io/SARS2-mut-fitness/mat_aa_fitness_correlations.html for the correlations among mutation effects estimated from the different sequence sets.

For the GISAID sequence set, we acknowledge the submitters of the sequences listed at the following URLs: https://doi.org/10.55876/gis8.230403ab, https://doi.org/10.55876/gis8.230403hg, https://doi.org/10.55876/gis8.230403ht, and https://doi.org/10.55876/gis8.230403tg.

### Counting mutations along the phylogenetic tree

We counted occurrences of each mutation in each viral clade using the UShER pre-built mutation-annotated tree [28, 29, 30] from May-11-2023 (http://hgdownload.soe.ucsc.edu/goldenPath/wuhCor1/UShER_SARS-CoV-2/2023/05/11/public-2023-05-11.all.masked.nextclade.pangolin.pb.gz), which contains all *∽*7-million SARS-CoV-2 sequences that are available in public databases. To make these counts at a per-clade level, we first subsetted the mutation-annotated tree on all sequences for each Nexstrain clade [69], retained only clades with at least 10^4^ sequences, and then used the matUtils program distributed with UShER to extract the nucleotide mutations on every branch of the each clade-subsetted mutation-annotated tree. For the analyses by geographic location (Figure 2), we subsetted on all sequences that began with “USA” or “England” as these were the two locations with the most publicly available sequences.

We then performed quality control by ignoring any branch that met any of the following criteria:

- it had more than four nucleotide mutations;
- it contained more than one nucleotide mutation that was a reversion to the Wuhan-Hu-1 reference sequence;
- it contained more than one nucleotide mutation that was a reversion to the founder sequence for that clade as provided at https://raw.githubusercontent.com/neherlab/SC2_variant_rates/7e738194a8c6592082f1caa9a6ca70cb68289790/data/clade_gts.json by [34];
- it contained more than one nucleotide mutation to the same codon.

The rationale for the first exclusion is that highly mutated branches are often indicative of sequencing errors of viral evolution in chronically infected humans, neither of which correspond to the pattern of typical SARS-CoV-2 transmission in acute infections. Because the virus’s evolution is very densely sampled, only a small fraction of branches have more than four mutations (Figure S10). The rationale for the second and third exclusions is that excess reversions can arise from base-calling pipelines that erroneously call low-coverage sites as reference. We ignore branches with multiple nucleotide mutations to the same codon (this is very rare) because as detailed below our method is only designed to make estimates for mutations that represent single-nucleotide changes from the clade founder. Note also that the mutation-annotated tree does not include insertion or deletion mutations, and so we only consider (and make estimates for) point mutations.

We then specified for exclusion certain mutations and sites that are prone to sequencing or base-calling errors. Specifically, we excluded

- the sites specified in Table S1 of [70] as being error prone;
- sites 5629, 6851, 7328, 28095, and 29362 since they had very high error rates in some clades;
- the problematic sites listed at https://github.com/W-L/ProblematicSites_SARS-CoV2, which are masked in the pre-built mutation-annotated tree;
- for each clade, the clade-specific sites listed in https://github.com/jbloomlab/SARS2-mut-fitness/blob/main/data/usher_masked_sites.yaml, which are masked in the pre-built mutation-annotated tree;
- for each clade, any mutation that was a reversion from the clade founder to the Wuhan-Hu-1 reference, and the reverse complements of these mutations.

The last exclusion criteria is because some bioinformatics pipelines called low-coverage sites as reference.

See https://github.com/jbloomlab/SARS2-mut-fitness/blob/main/results/mutation_counts/aggregated.csv for the final counts of each nucleotide mutation in each clade; note that this file also contains excluded mutations.

### Calculation of expected counts

To calculate the expected counts for each nucleotide mutation, we analyzed just the four-fold degenerate sites in each clade in an approach paralleling that of [31]. Specifically, we identify all non-excluded four-fold degenerate sites in each clade founder. We then count nucleotide mutations just at those sites in each clade, and calculate the expected per-site number of mutations from nucleotide *x* to *y* as the total number of *x* to *y* mutations at four-fold degenerate sites divided by the number of four-fold degenerate sites with *x* as the parental identity. This analysis is done at the clade level for two reasons: referencing mutations to the clade founder (rather than the Wuhan-Hu-1 reference) limits problem with the approach that would arise at sites that substitute multiple times in the history of a sequence (since each clade is a relatively high-identity group multiple mutations at the same site within a clade are very rare), and because it is know that SARS-CoV-2 mutation rates vary somewhat among clades [31, 32]. We only retain clades with at least 5000 mutations at four-fold degenerate sites in order to avoid inaccurate estimates of expected counts due to low sampling of mutations.

### Mutational effects from actual versus expected counts

To estimate the effects of mutations, we simply compare the expected counts of each nucleotide mutation to the actual counts in the pre-built mutation-annotated tree. See https://github.com/jbloomlab/SARS2-mut-fitness/blob/main/results/expected_vs_actual_mut_counts/expected_vs_actual_mut_counts.csv for these expected versus actual counts on a per-clade basis; note that this file also includes counts at excluded sites. To estimate the effects of mutations, we first sum the counts of all non-excluded nucleotide mutations that encode each amino-acid mutation to convert the nucleotide counts to amino-acid counts. In doing this, we exclude any mutations that are not from the clade-founder codon identity: in other words, we ignore sequences with histories that involve multiple mutations at the same codon in the same clade (this is a caveat of the approach, although because each clade is relatively high identity it does not have a major effect). For the overall estimates reported in this paper, we also sum these counts across all retained clades; for the analyses in Figure 2 we also make estimates without summing across clades and only for counts from sequences from specific geographic locations. We then compute the estimated fitness Δ *f* of each mutation as simply the natural logarithm of the ratio of actual to expected counts after adding a pseudocount of *P −* 0.5 to each count, namely 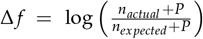

Note that these mutation-effect estimates will have more statistical noise the smaller the value of the expected counts for each mutation. Therefore, we also track the expected counts alongside the estimates. In this paper, we only show estimates for mutations with expected counts of at least 10 unless otherwise noted. However, the figures link to interactive legends that allow adjustment of this threshold: larger values (eg, 20 or more) will lead to slightly more accurate estimates but drop some mutations, lower values can be used if you need a noisier estimate for a mutation that has less then 10 expected counts.

See https://github.com/jbloomlab/SARS2-mut-fitness/blob/main/results/aa_fitness/aamut_fitness_all.csv for the estimates of amino-acid mutation effects across all clades, and see https://github.com/jbloomlab/SARS2-mut-fitness/blob/main/results/aa_fitness/aamut_fitness_by_clade.csv for the clade-specific estimates. The all-clade estimates of mutation effects are what are shown in Figure 3.

For the clade correlations plotting in Figure 2, we only include clades with at least 5 *×* 10^5^ expected counts across all sites, as only these clades have enough counts for reasonable per-clade estimates.

### Mutation effects to amino-acid fitnesses

For the final estimates of amino-acid fitnesses shown in the heatmaps such as in Figure 5, we need a single estimate for each amino acid. This is straightforward for sites that have the same amino-acid identity in all clade founders: the “wildtype” residue shared across all clades has a fitness of zero, and all other amino acids have fitnesses equal to the effect of mutating from the “wildtype” to that amino acid. However, for sites that change amino-acid identity between clade founders, things are more complicated and we need to take the extra step below.

For each clade have estimated the change in fitness Δ *f*_*xy*_ caused by mutating a site from amino-acid *x* to *y*, where *x* is the amino acid in the clade founder sequence. For each such mutation, we also have *n*_*xy*_ which is the number of expected mutations from the clade founder amino acid *x* to *y*. These *n*_*xy*_ values are important because they give some estimate of our “confidence” in the Δ *f*_*xy*_ values: if a mutation has high expected counts (large *n*_*xy*_) then we can estimate the change in fitness caused by the mutation more accurately, and if *n*_*xy*_ is small then the estimate will be much noisier.

However, we would like to aggregate the data across multiple clades to estimate amino-acid fitness values at a site under the assumption that these are constant across clades. Things get complicated if not all clade founders have the same amino acid identity at a site. For instance, let’s say at our site of interest, the clade founder amino acid is *x* in one clade and *z* in another clade. For each clade we then have a set of Δ *f*_*xy*_ and *n*_*xy*_ values for the first clade (where *y* ranges over the 20 amino acids, including stop codon, that aren’t *x*), and another set of up to 20 Δ *f*_*zy*_ and *n*_*zy*_ values for the second clade (where *y* ranges over the 20 amino acids that aren’t *z*).

From these sets of mutation fitness changes, we’d like to estimate the fitness *f*_*x*_ of each amino acid *x*, where the *f*_*x*_ values satisfy Δ *f*_*xy*_ = *f*_*y*_ *− f*_*x*_ (in other words, a higher *f*_*x*_ means higher fitness of that amino acid). When there are multiple clades with different founder amino acids at the site, there is no guarantee that we can find *f*_*x*_ values that precisely satisfy the above equation since there are more Δ *f*_*xy*_ values than *f*_*x*_ values and the Δ *f*_*xy*_ values may have noise (and is some cases even real shifts among clades due to epistasis). Nonetheless, we can try to find the *f*_*x*_ values that come closest to satisfying the above equation.

First, we choose one amino acid to have a fitness value of zero, since the scale of the *f*_*x*_ values is arbitrary and there are really only 20 unique parameters among the 21 *f*_*x*_ values (there are 21 amino acids since we consider stops, but we only measure differences among them, not absolute values). Typically if there was just one clade, we would set the wildtype value of *f*_*x*_ = 0 and then for mutations to all other amino acids *y* we would simply have *f*_*y*_ = Δ *f*_*xy*_. However, when there are multiple clades with different founder amino acids, there is no longer a well defined “wildtype”. So we choose the most common non-stop parental amino-acid for the observed mutations and set that to zero. In other words, we find *x* that maximizes ∑_*y*_ *n*_*xy*_ and set that *f*_*x*_ value to zero.

Next, we choose the *f*_*x*_ values that most closely match the measured mutation effects, weighting more strongly mutation effects with higher expected counts (since these should be more accurate). Specifically, we define a loss function as

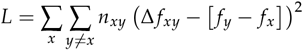

where we ignore effects of synonymous mutations (the *x* ≠ *y* term in second summand) because we are only examining protein-level effects. We then use numerical optimization to find the *f*_*x*_ values that minimize that loss *L*.

Finally, we would still like to report an equivalent of the *n*_*xy*_ values for the Δ *f*_*xy*_ values that give us some sense of how accurately we have estimated the fitness *f*_*x*_ of each amino acid. To do that, we tabulate *N*_*x*_ = ∑_*y*_ (*n*_*xy*_ + *n*_*yx*_)as the total number of mutations either from or to amino-acid *x* as the “count” for the amino acid. Amino acids with larger values of *N*_*x*_ should have more accurate estimates of *f*_*x*_.

See https://github.com/jbloomlab/SARS2-mut-fitness/blob/main/results/aa_fitness/aa_fitness.csv for these overall amino-acid fitness estimates.

### Site numbering and protein naming

All sites are numbered according to the sequential Wuhan-Hu-1 reference numbering scheme, using the reference sequence at http://hgdownload.soe.ucsc.edu/goldenPath/wuhCor1/bigZips/wuhCor1.fa.gz. The protein annotations are taken from the associated GTF at http://hgdownload.soe.ucsc.edu/goldenPath/wuhCor1/bigZips/genes/ncbiGenes.gtf.gz. Those protein annotations refer to the polyproteins encoding the non-structural proteins as ORF1a and ORF1ab. To convert to from ORF1ab numbering/naming to the nsp-based naming (eg, nsp1, nsp2, etc) we use the conversions specified under “orf1ab_to_nsps” in https://github.com/jbloomlab/SARS2-mut-fitness/blob/main/config.yaml, which are in turn taken from Theo Sanderson’s annotations at https://github.com/theosanderson/Codon2Nucleotide/blob/main/src/App.js.

### Comparison to deep mutational scanning

Deep mutational scanning data were taken from published studies [9, 49, 21, 50, 22], using the data at the links specified under the “dms_-datasets” key in https://github.com/jbloomlab/SARS2-mut-fitness/blob/main/config.yaml. For the spike deep mutational scanning [9] we only included mutations with “times seen” values of at least three in the deep mutational scanning. The RBD data [49] include measurements for two phenotypes (ACE2 affinity and RBD expression), and one of the Mpro studies [21] includes measurements for three different phenotypes in yeast (growth, FRET, and transcription factor activity). Figures 4, S5, and S6 shows the effect averaged across all phenotypes measured by each of these studies. For plots that break the correlations out by phenotype, see https://jbloomlab.github.io/SARS2-mut-fitness/dms_S_all_corr.html and https://jbloomlab.github.io/SARS2-mut-fitness/dms_nsp5_all_corr.html.

### Comparison to dN/dS and other mutation-effect prediction algorithms

For the comparison to the dN/dS approaches shown in Figure S5, we used the dN/dS values available at https://github.com/spond/SARS-CoV-2-variation [71] for all SARS-CoV-2 sequences, which were calculated using the FEL approach [53]. The dN/dS ratios only provide a single number for each site, which cannot be directly compared to either the mutation-effect estimates or the deep mutational scanning, which estimate the effects of individual amino-acid mutations. We therefore computed site-summary metrics of the mutation-effect estimates and the deep mutational scanning as the average effect of all measured amino-acid mutations at each site, excluding stop codons. The correlations in Figure S5 are with those site-summary metrics.

We also compared both our mutation-effect estimates and the spike deep mutational scanning measurements [9] to predictions from three other algorithms:

- the EpiScores reported by Maher et al (2022) [25],
- the DCA mutability scores reported by Rodriguez-Rivas et al (2022) [26], and
- the EVE scores reported by Thadani et al (2023) [27].

These comparisons are shown in Figure S6. The Maher et al (2022) and Thadani et al (2023) studies report mutation-level predictions and so are compared directly to the deep mutational scanning our mutation-effect estimates; Rodriguez-Rivas et al (2023) report only site-level metrics and so are compared to site-summary metrics as for the dN/dS analysis.

### Derivation of relationship between actual to expected count ratio and viral fitness

The ratio of actual to expected counts that we calculate in this paper is related to the probability that we observe a viral lineage containing an occurrence of a specific mutation among sequenced human SARS-CoV-2. This probability depends on three factors: the fitness effect of the mutation, the fraction of all SARS-CoV-2 viruses that are sequenced (sampling intensity), and the growth dynamics of the viral population. In the supplementary appendix, we derive the approximate relationship between this probability as a function of the fitness cost *s* and sampling intensity *ϵ* for deleterious mutations for both a constant and exponentially growing viral population.

We show that for a constant viral population size, the probability of observing a lineage containing a deleterious mutation with cost *s* is roughly 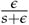when *s*^2^ *> ∈*, and more weakly dependent on *s* for smaller fitness costs (when *s*^2^ < *ϵ*). The intuitive explanation is that the average size of a mutant lineage with fitness cost *s* is 1/*s* and we basically ask whether we sample the lineage before it disappears. If we sample more intensely (larger *ϵ*), whether a lineage gets sampled depends primarily on the stochastic dynamics and little on the fitness effect. With a typical sampling intensity for SARS-CoV-2 between 1/1000 and 1/100, this means our approach is sensitive to fitness effects larger than a few percent per serial interval; mutations with fitness costs smaller than that will not show an appreciable difference from neutral mutations in their ratio of actual to expected accounts.

In an exponentially growing population, the probability of observing a mutant lineage with fitness cost *s* again scales as *∽*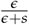 if *sT >* 1, where *T* is the time over which the variant has expanded. If *T* is *∽* months, that is 20 generations, which again corresponds to *s* of at least a few percent for *sT >* 1. For mutations with smaller fitness costs, the dependence scales more as ~ *ϵ* (1 *− sT*).

Overall, these calculations indicate that for multiple different growth dynamics of the viral population, the ratio of expected to actual counts will scale inversely with the fitness cost of deleterious mutations for mutations with costs that exceed a few percent. Note that the approach we use in this paper does not account for variation in sampling intensity across space or time, does not attempt to adjust for changes in viral growth dynamics over time, uses the heuristic formula of calculating the effect as the log ratio of counts, and applies this same formula to all mutations regardless of whether they are deleterious, neutral, or beneficial. A more complete derivation might try to calculate the fitness effects from the full distribution of lineage sizes more rigorously and incorporate information about the sampling intensity and viral growth dynamics. However, such a derivation (if possible at all) is beyond the scope of this study, and we also note that good empirical data is generally lacking to precisely account for sampling intensity and viral growth dynamics over the full span of time and space from which the sequences we analyze are drawn. The key point of the derivations for our current study is simply that our approach should be sensitive to detecting the effects of mutations with fitness costs greater than a few percent.

## Acknowledgments

We thank Angie Hinrichs for providing the pre-built mutation-annotated trees and thank the UShER team for promptly answering and addressing GitHub issues related to use of this package and its pre-built trees. We thank the sequence submitters to GISAID, who are listed in the tables cited in the Methods section. The work of JDB was supported in part by the NIH/NIAID under grants U19AI171399 and R01AI141707, and contract 75N93021C00015. JDB is an Investigator of the Howard Hughes Medical Institute.

## Supplementary appendix deriving relationship between fitness cost and ratio of expected to actual counts

With millions of SARS-CoV-2 sequences shared publicly, almost all mutations that are tolerated by the virus are observed dozens to hundreds of times. Where on the tree and how often on the tree we observe specific mutations has information about the effects of these mutations on viral spread. The mutation rate depends on the nucleotides involved and possibly on the sequence context and other viral determinants, but for the purpose of this derivation, we will assume the neutral rate *μ* is known. If the mutation is neutral, the total number of times the mutation is observed on the tree is *μT*, where *T* is the total length of the tree (assuming that the mutation never reached high frequency which is true for almost all mutations, particularly when mutations are counted on a per-clade basis relative too the clade founder as done above). If a mutation reduces fitness, the lineages descending from branches on which this mutation happened will spread more slowly than those without this mutation. As a result, the down-stream subclades are smaller and more short lived, which in turn means that they will be less likely to be sampled and represented in the tree. To infer a mutation’s effect on fitness, we need to calculate how the probability of observation depends on this fitness effect.

For a mutation to be represented in the tree, one of its descendants has to be sampled and sequenced. If the total number of descendants is *w* and the sampling fraction is *ϵ*, the probability that the mutation is present in the tree is

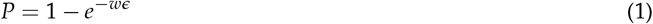

*W* is a random number that depends on the realization of the transmission process, which is commonly modeled by a branching process with birth rate *b* and death rate *d*. The death rate here corresponds to clearing an infection, the birth rate to onward transmission. The latter is affected by the fitness cost of the mutation.

To obtain insight how the probability of observing a lineage depends on parameters, we calculate the probability *p*(*w, T t*) that a lineage had an integrated size 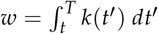, where *t* is the birth time of the lineage, *T* is the current time, and *k*(*t*′) is the size of the lineage at time t′. To calculate *p*(*w, T k*), we generalize it slightly to *p*(*W, T k, t*), where *k* is the number of individuals at the start time *t*. This quantity obeys the following “first-step” equation:

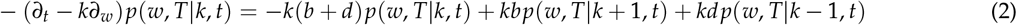

We will solve for the Laplace transform 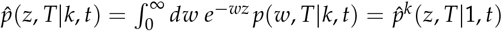. Using the following identity for the derivative of the Laplace transform

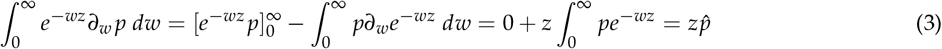

and setting *k* = 1, we have

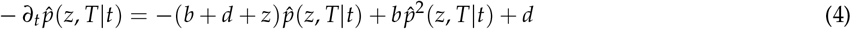

This simplifies further to if we substitute 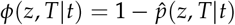.

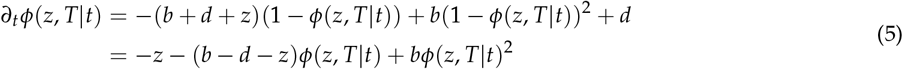

where it is important to note that the derivative is with respect to the first time point and the interval *T − t* is shrinking with increasing *t*.

### Constant birth and death rate

If the fitness effect of the mutation in question is detrimental and the overall population is constant (background *b*_0_ = *d*_0_), all mutant lineages will eventually die out and we can consider large *T − t* and the long time asymptotic *∂tϕ*(*z, T/ t*) = 0. Further define *b* = *b*_0_ *− s* and *d* = *d*_0_ where *s* is the fitness cost of the mutation (so larger values indicate a greater fitness cost). The steady state generating function is then

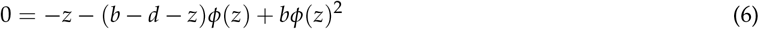

with solution

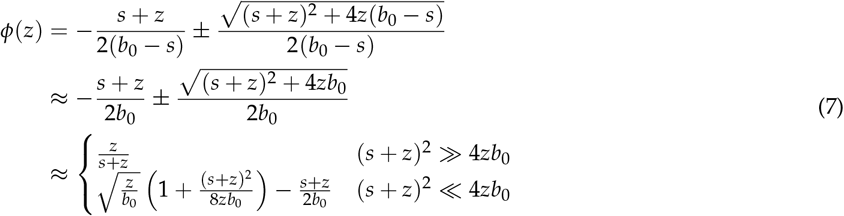

Since *ϕ*(*z*) = 1 *e*^*−wz*^ *p*(*w*) *dw, ϕ*(*∈*) is exactly the probability that a lineage is sampled when the entire population is sampled at rate *ϵ*. We thus expect two regimes: if the square of the fitness cost exceeds the sampling intensity (typically at 1% or less), the probability of sampling a lineage is essentially inversely proportional to the fitness cost. The sampling probability of lineages with smaller costs effects depends less strongly on *s*. Their sampling mostly comes down to stochasticity independent of the fitness cost.

### Growing populations

In many scenarios relevant for lineages that arise during a viral outbreak, the background population isn’t constant but is undergoing a rapid exponential expansion. The background birth rate *b*_0_ is bigger than *d*_0_ in this case. Since the population is growing, deleterious mutations can increase in frequency deterministically and we can not send the *t* to infinity as before. Instead, we need to integrate

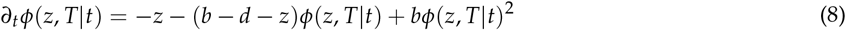

backwards in time starting from *ϕ*(*z, T T*) = 0 at *t* = *T*. While *ϕ*(*z, T t*) is small and the quadratic term can be neglected, this is approximately solved by

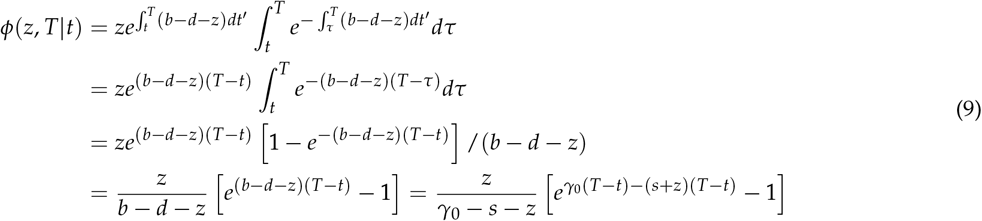

where *γ*_0_ is the growth rate of the background population.

At longer times when *ze*^*γ*^_0_ ^(*T−t*)^ *∽* 1 and *ϕ* is no longer small, *ϕ* tends towards a constant value determined by the same quadratic equation as above. This limit is neither interesting or relevant for the present purpose, since there are very few lineages that emerged early enough to have saturated *ϕ*. Instead, we need to average *ϕ* (the linear approximation) over all the time points when the lineage could have arisen.

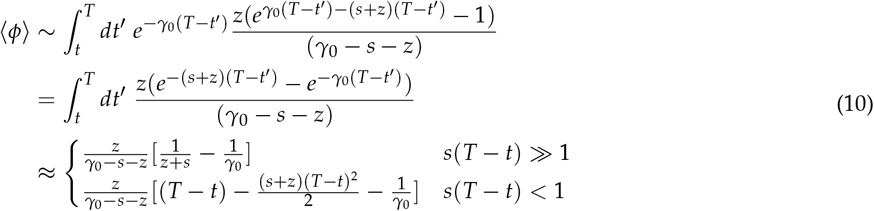

This derivation assumed that *γ*_0_(*T − t*) ≫ 1, i.e. that the overall population size has expanded substantially. The most relevant fitness effects will be those with *s*(*T − t*) *>* 1, that is the fitness effect has strong effect on variant frequency, but *s* < *γ*_0_ such that the variant is still spreading and can give rise to large lineages in an expanding variant. In this case, the above simplifies to

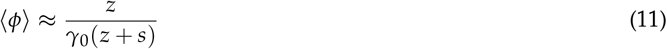

In a variant that has been growing with rate *γ*_0_ for a time *τ* = *T − t* and sampled with *z* = *ϵ*, we thus expect that the number of times we observe separate mutant lineages depends on *s* as

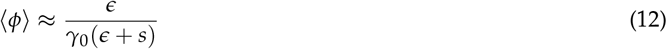

This has a very similar behavior as the solution for constant population size, which suggests that the overall dependence on *s* is robust and we can assume that the number of times a mutation is observed is inversely proportional to its effect on fitness. The same basic dependency is observed at steady state in a quasi-species [65]. In a constant population, this relationship breaks down for dense sampling *∈ >*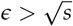. In growing population, the approximation fails if the product of fitness effect and the time over which the variant has grown, *sτ*, is small, i.e., if the fitness cost does not affect variant frequency strongly. In these cases, there is still a dependence on *s*, but it is weaker.

## Supplementary figures

**Figure S1.**
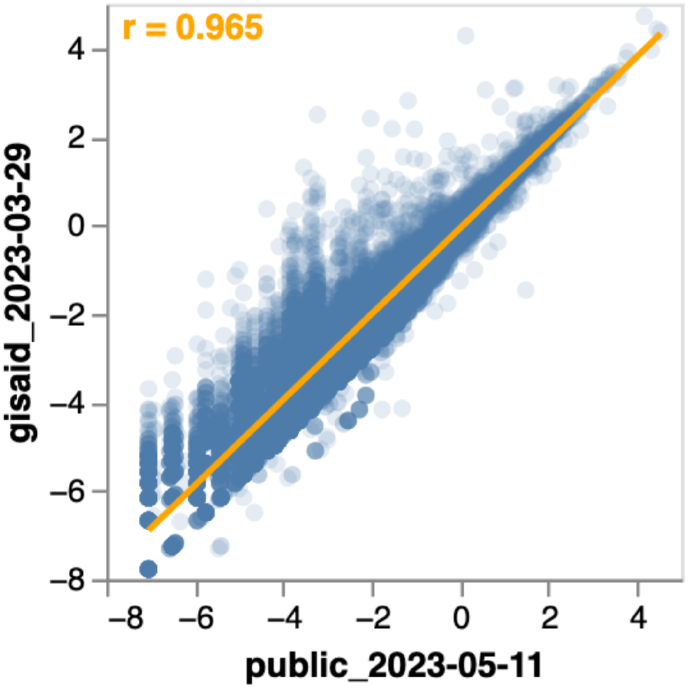
Correlation between amino-acid fitness estimates from the *∽*7-million sequence set of all publicly available sequences as of May-11-2023, or the set of all ~14-million sequences in GISAID as of March-29-2023. The orange text in the upper-left corner of the plot shows the Pearson correlation. The high correlation indicates that our estimates are not substantially limited by noise related to statistical sampling of mutation counts, since using another tree with twice as many sequences does not substantially alter the estimates. Note that the main text figures all use the smaller*∽*7-million publicly available sequence set. See https://jbloomlab.github.io/SARS2-mut-fitness/mat_aa_fitness_correlations.html for an interactive version of this plot that also shows the correlations for other sets of publicly available sequences.

**Figure S2.**
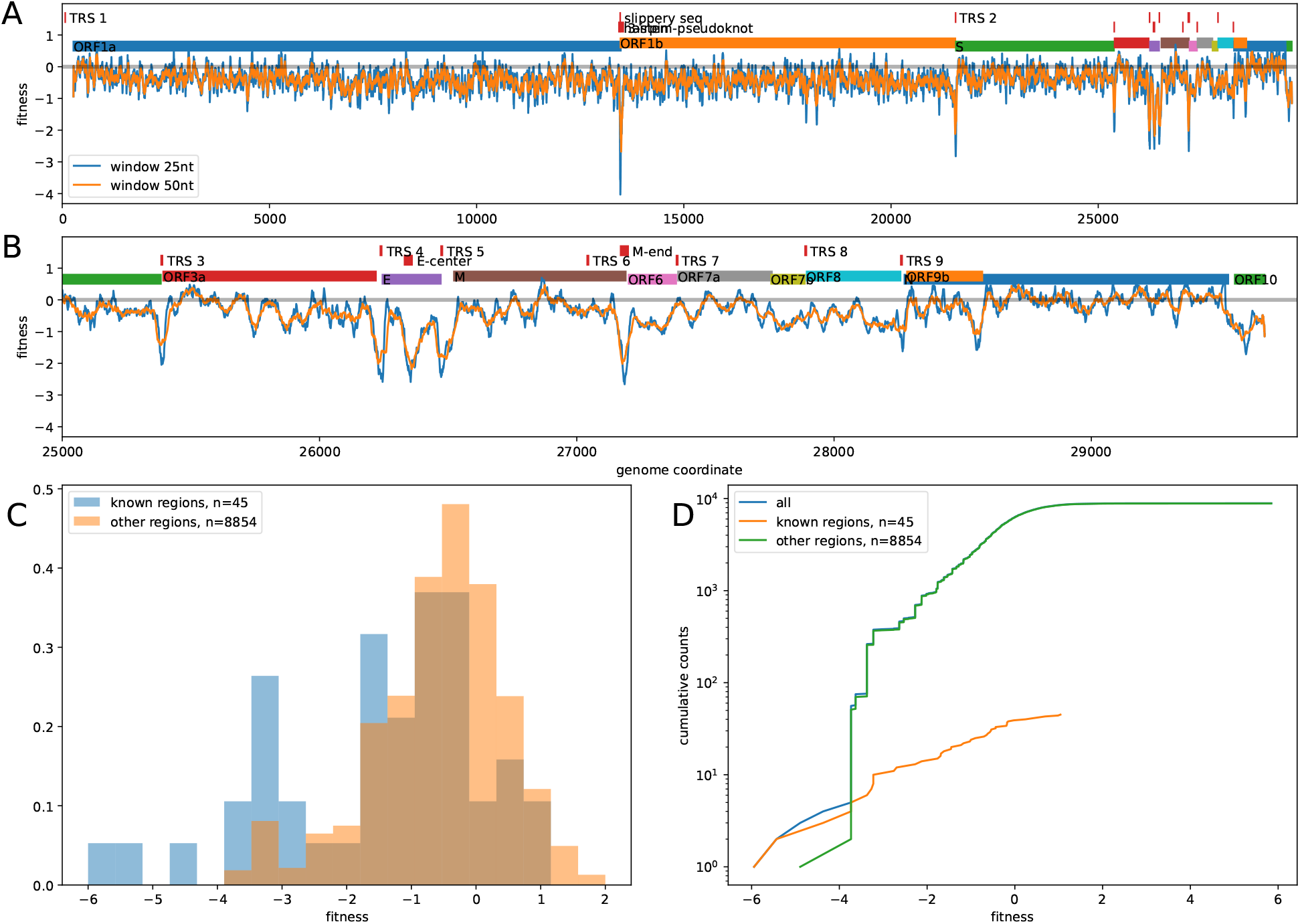
Non-coding constraint is concentrated in a few isolated regions in SARS-CoV-2 genome. **(A), (B)** The average fitness cost of synonymous mutations (do not change amino acids) in ORF1ab, S, E, M, N, and ORF3a, non-coding regions between ORFs, and all positions in ORF6, ORF7a, ORF7b, ORF8, ORF10 (as these ORFs tolerate stop codons, see Figure 3B). Panel A shows the entire genome, panel B zooms into the 3’ end of the genome. Fitness effects are averaged in sliding windows of size 25 and 50. With the exception of a small number of well defined regions, in most of the genome these averages fluctuate around zero. Regions with large fitness costs for non-coding mutations correspond to the ribosomal slippage site at the end of ORF1a, the transcription regulatory sites of S, ORF3a, E, M, and N, as well as two regions in the center of E and the end of M (see red bars in the top part of each panel) [72]. Notably, TRS sites of ORF6, ORF7a and ORF8 don’t show strong signal of conservation, consistent with their tolerance of stop codons. **(C), (D)** The distribution of inferred fitness cost of four-fold degenerate sites at regions of known non-coding constraint within the protein coding genes (the ribosomal slippage and TRS sites) versus all other four-fold degenerate sites. In total, 25 four-fold degenerate sites are in these regions of known constraint, whereas 4844 four-fold degenerate sites are outside these regions. Panel C shows that a large fraction of mutations in these known regions of constraint are deleterious, while the distribution for the remaining four-fold degenerate sites is centered around 0 between -2 and +1. Furthermore, strongly deleterious mutations with fitness estimate below -3 predominantly fall in the 25 sites of known non-coding constraint. If the two additional conserved regions in E and M are included, all but one mutation outside these regions has a fitness cost estimate above -3.

**Figure S3.**
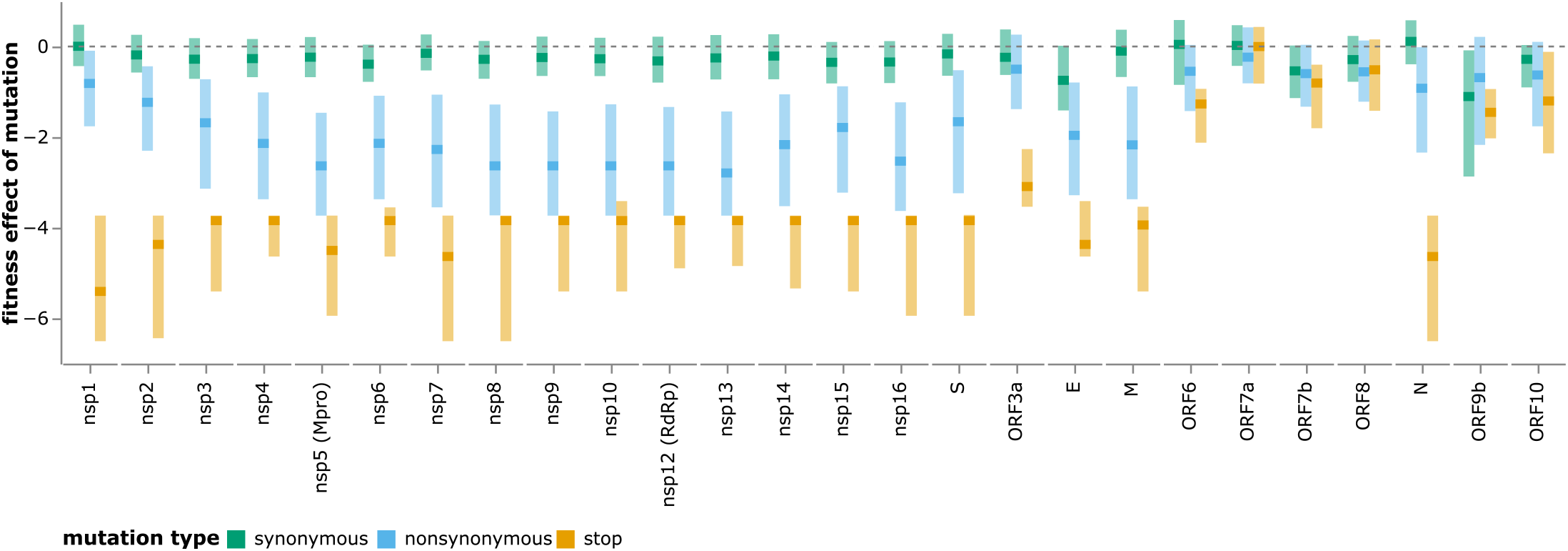
A version of Figure 3B but with genes ordered by position in the genome rather than extent of constraint on nonsynonymous mutations. See https://jbloomlab.github.io/SARS2-mut-fitness/effects_dist_position_order.html for an interactive version of this plot.

**Figure S4.**
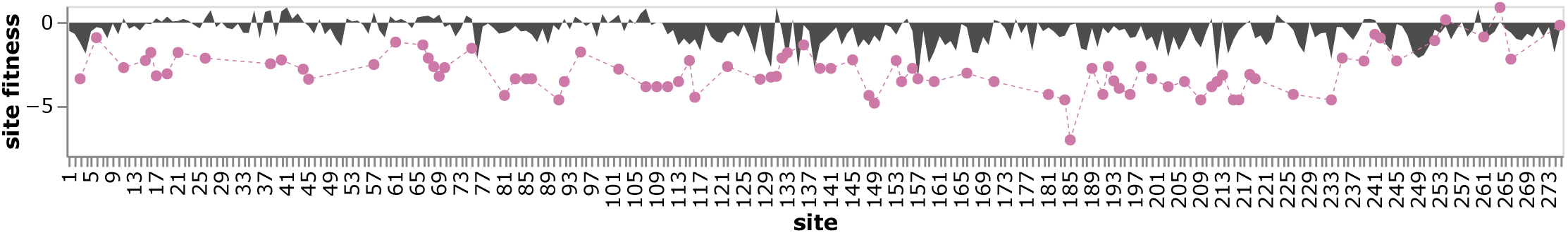
Effects of stop-codon and amino-acid mutations across ORF3a. The black area plot shows the mean effect of all amino-acid mutations at each site, and the purple points show the effects of stop codon mutations. There is strong selection against stop codons (negative effects) for all but the C-terminus of ORF3a, but only a few positions show strong selection against amino-acid substitutions. See https://jbloomlab.github.io/SARS2-mut-fitness/ORF3a.html for an interactive version of this plot along with zoomable heatmap of the effects of specific amino-acid substitutions.

**Figure S5.**
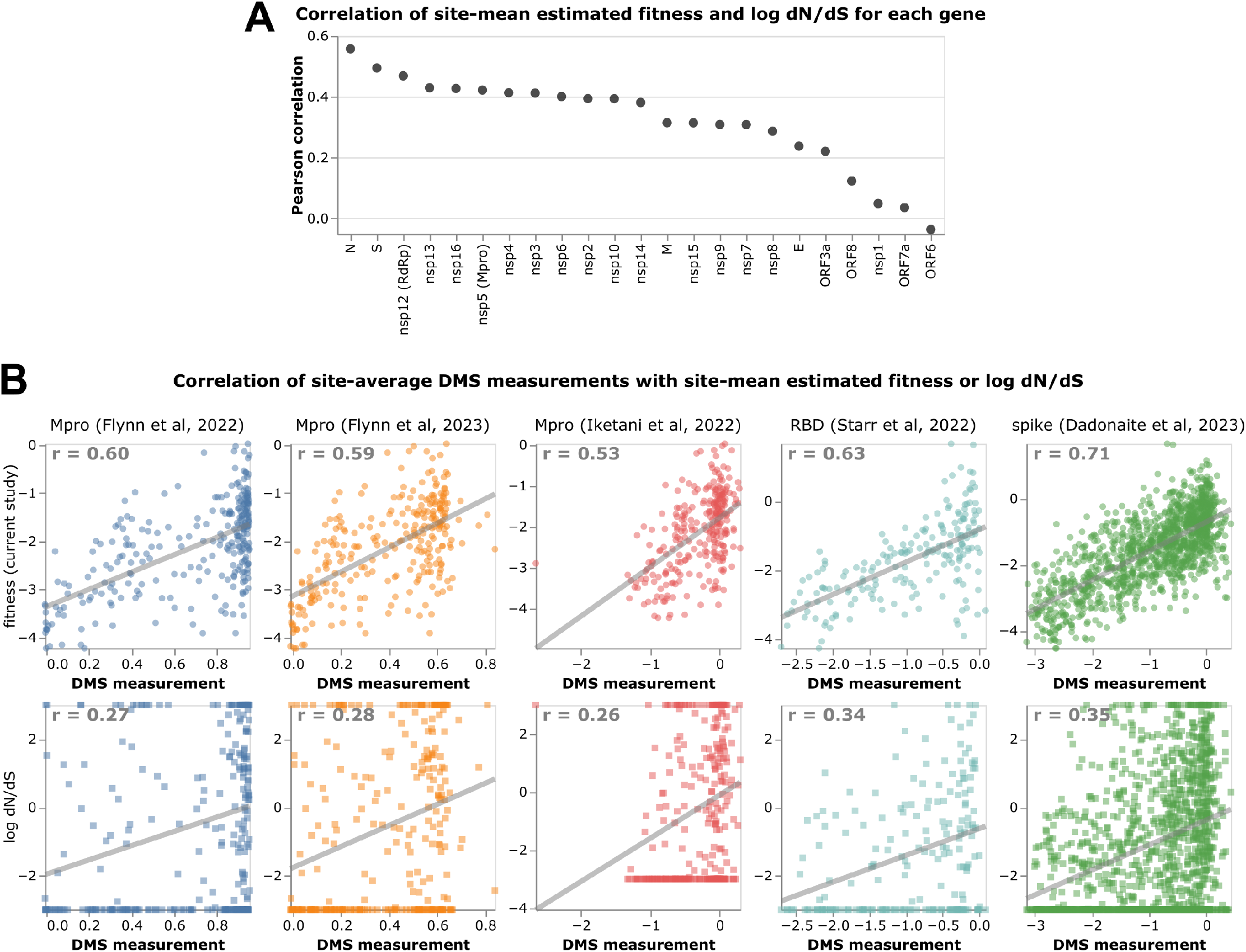
The site-mean of the estimated fitness effects outperforms dN/dS ratios as an indicator of functional constraint as assessed by correlation with deep mutational scanning measurements. **(A)** Pearson correlation of log dN/dS ratio at each site with the mean estimated fitness of all amino acid mutations at each site from the current study. For most genes, the log dN/dS ratio is correlated with the mean fitness effect, but the correlations are only modest (always substantially less than one). The exception is the accessory proteins which are mostly under minimal amino-acid level selection (Figure 3B) and only show a weak correlation. **(B)** Correlation of the log dN/dS ratio or the mean estimated fitness effect at each site with the mean impact of amino-acid mutations at that site as experimentally measured by deep mutational scanning. The numbers in the upper right of each panel give the Pearson correlation. In every case, the mean estimated fitness effect (top row) is much better correlated with the experimental measurements than the log dN/dS (bottom row). The dN/dS ratios are taken from https://github.com/spond/SARS-CoV-2-variation. See https://jbloomlab.github.io/SARS2-mut-fitness/dnds_corr.html for an interactive version of this plot.

**Figure S6.**
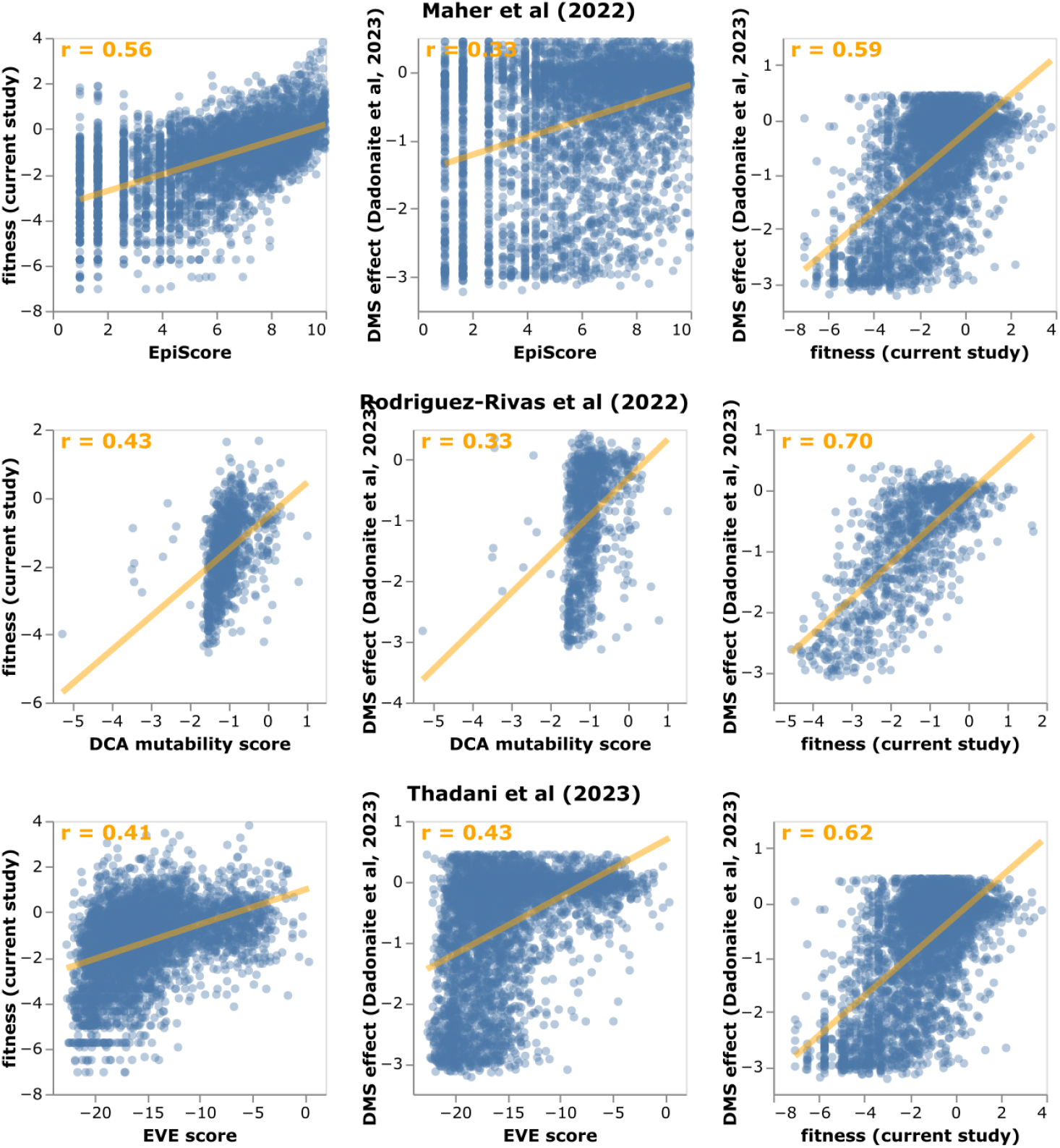
The mutation effect estimates from the current study are substantially better correlated with deep mutational scanning measurements for spike than predicted mutation effects made using three other approaches. Each row shows the correlation of the fitness estimates from the current study and full-spike deep mutational scanning measurements versus values from another study. The Maher et al (2022) and Thadani et (2023) studies make predictions for individual amino-acids, so those correlations are with the mutation-level fitness estimates and deep mutational scanning measurements. The Rodriguez-Rivas et al (2022) study only reports site-level predictions, so those correlations are with the site mean fitness estimates and deep mutational scanning measurements. Each row only shows mutations or sites with a reported value from all three approaches. The numbers in the upper right give the Pearson correlation. See https://jbloomlab.github.io/SARS2-mut-fitness/comparator_corr.html for an interactive version of this plot.

**Figure S7.**
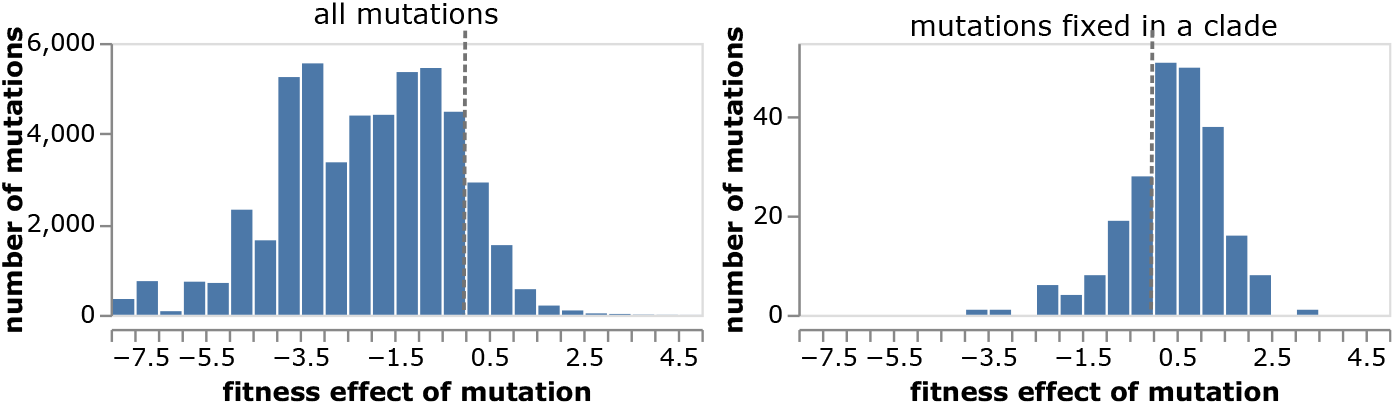
Distribution of fitness effects of all amino-acid mutations relative to Wuhan-Hu-1, and just those mutations that fixed in at least one clade of SARS-CoV-2 (using the Nextstrain clade definitions). The vertical dashed line at zero indicates the effect of a neutral mutation. See https://jbloomlab.github.io/SARS2-mut-fitness/clade_fixed_muts_hist.html for an interactive version of this plot that allows adjustment of the minimum expected count threshold.

**Figure S8.**
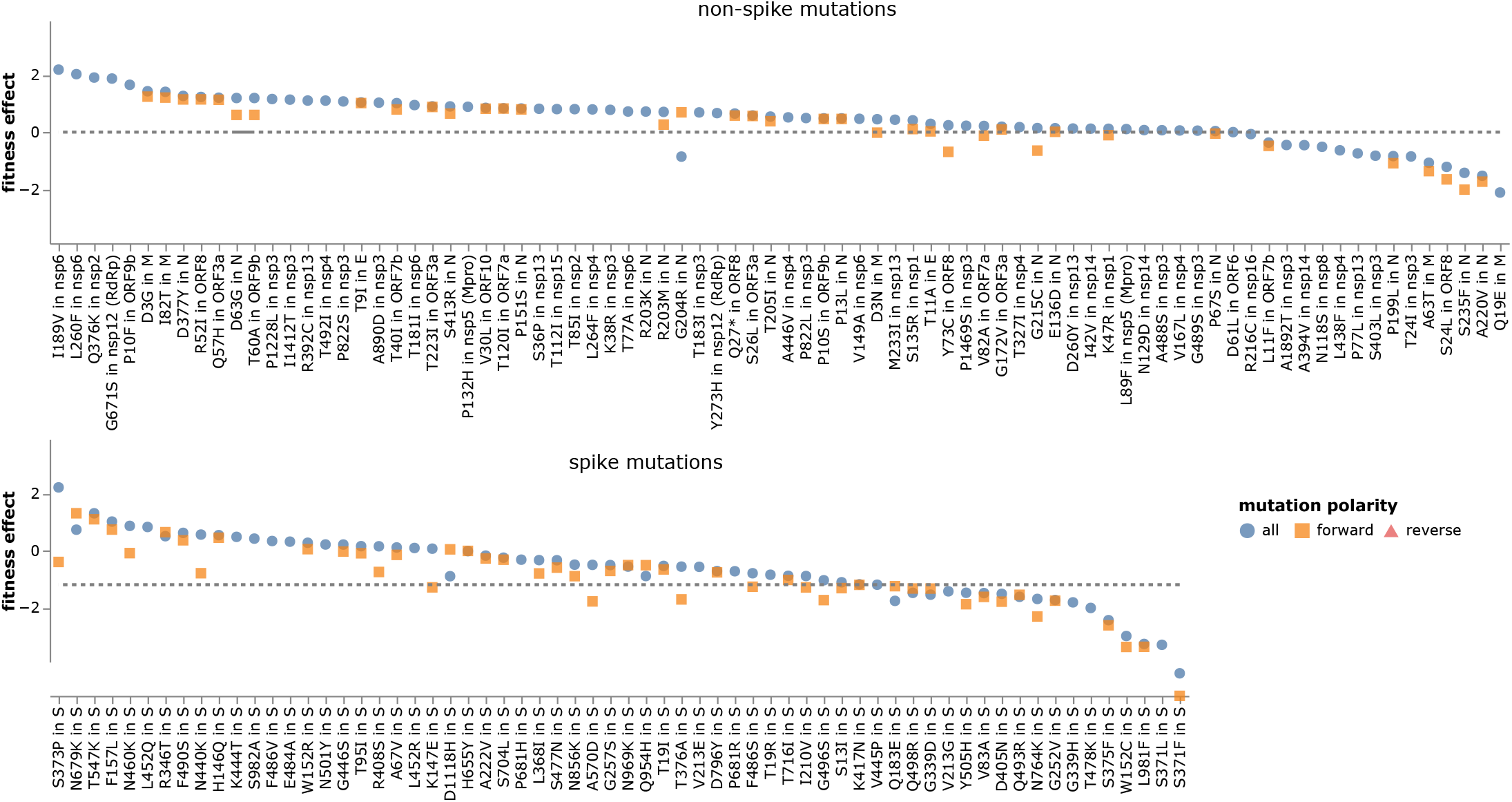
Effects of individual mutations that fixed in at least one clade of SARS-CoV-2, faceted by whether they are in spike or another protein. “Mutation polarity” indicates if the point shows the effect of the mutation estimated using all viral clades (including those that have fixed the mutation), or just from direct forward occurrences of the mutation in clades in which it has not yet fixed. Some mutations are estimated to be more favorable when including clades in which they have fixed (blue circles) in addition to just clades in which it has not yet fixed (orange squares)—when this occurs, it suggests epistatic entrenchment of the mutations [38, 60]. Note that clades in which a mutation has already fixed contribute to estimates of its fitness via estimates of the effect of its reversion and via estimates of the effects of mutations to other amino acids at the same site. See https://jbloomlab.github.io/SARS2-mut-fitness/clade_fixed_muts.html for an interactive version of this plot.

**Figure S9.**
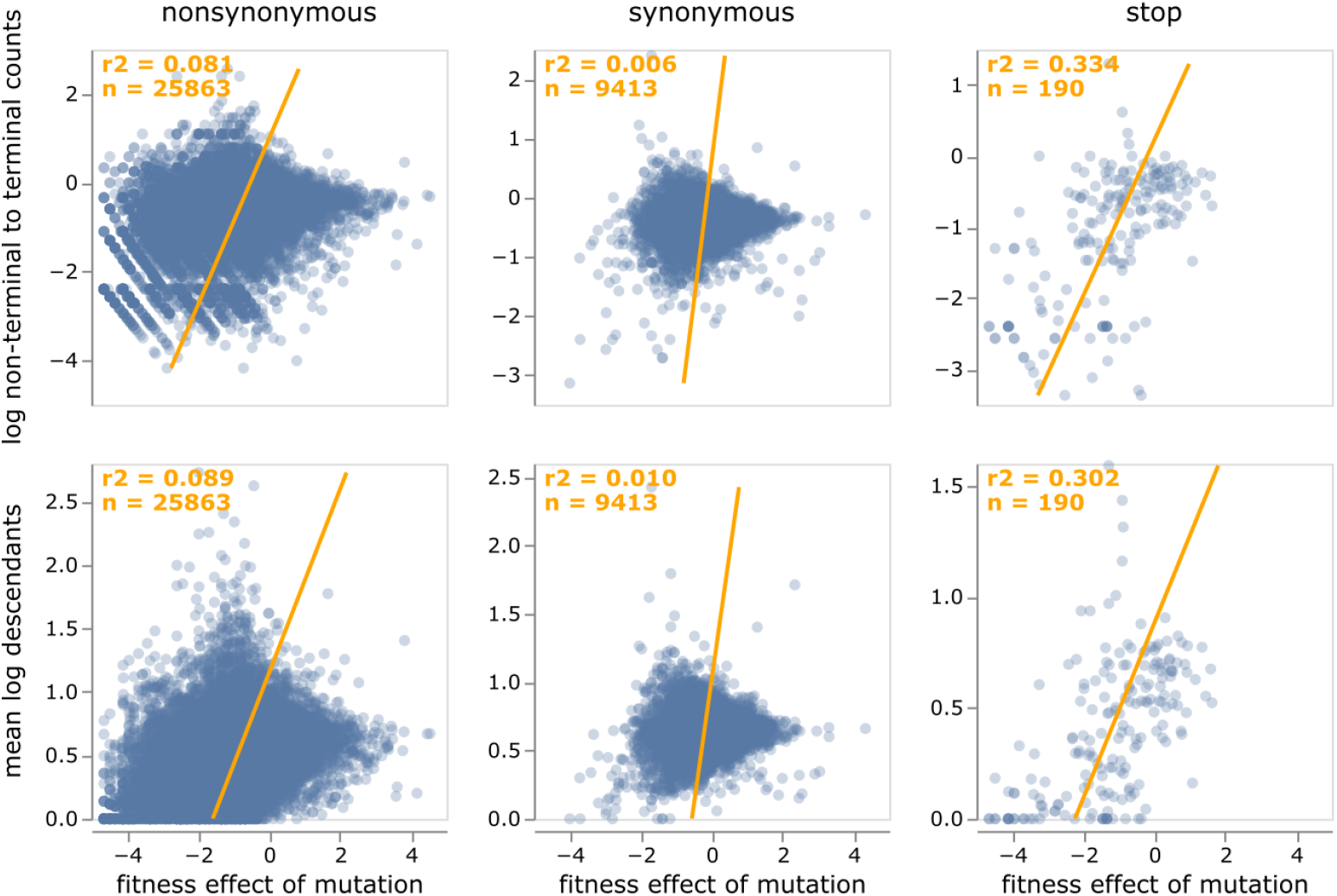
Relationship between fitness effects of mutations and two measures of the number of descendants. At top is shown the log ratio of counts of the mutation on non-terminal (internal) to terminal (tip) branches; larger values indicate mutations more likely to be found in viruses that leave descendants. At bottom is shown the mean log number of tip descendants that share all the mutations on each branch containing the mutation of interest; larger values again indicate mutations more likely to be found in viruses that leave more descendants. Each point is an amino-acid mutation, the orange line is a least-squares regression, and the orange text in the upper left give the number of mutations and the Pearson correlation coefficient. This plot shows only mutations with at least 10 expected counts and 5 actual counts. See https://jbloomlab.github.io/SARS2-mut-fitness/fitness_vs_terminal.html for an interactive version of this plot that allows filtering by the number of actual or expected counts, or by gene. The number of descendants is calculated using the “leaves_sharing_mutations” variable of the UShER mutation-annotated tree.

**Figure S10.**
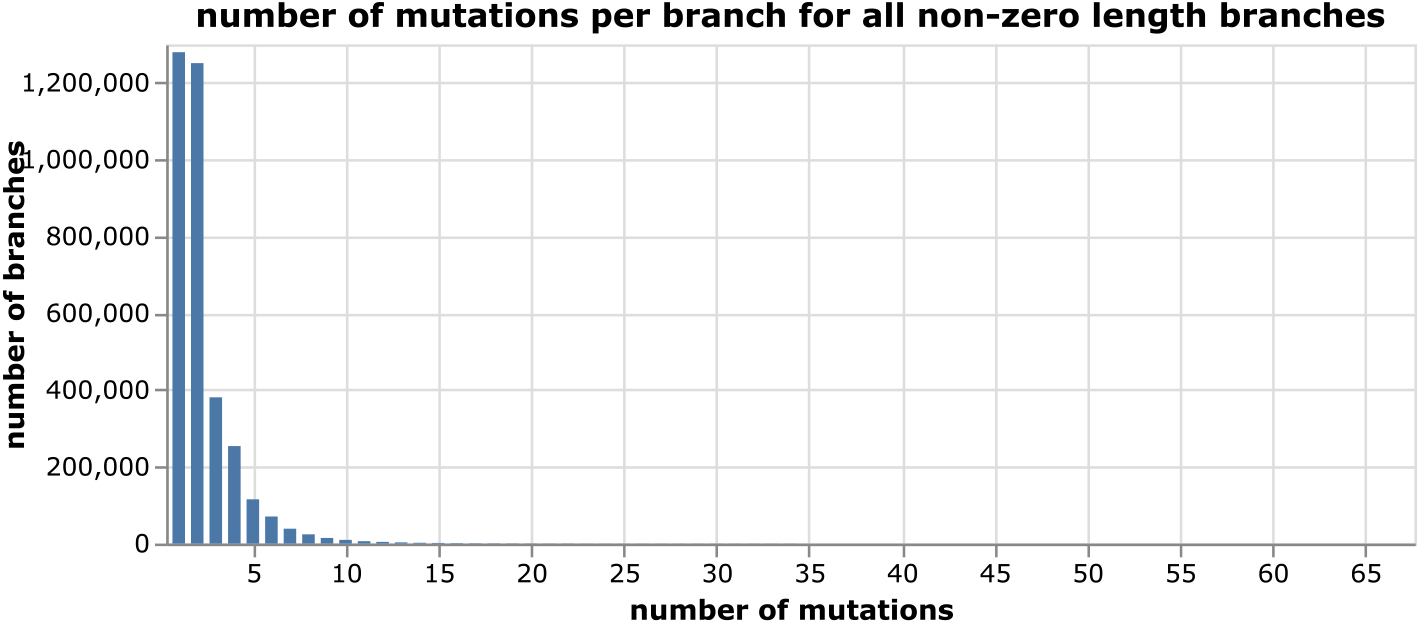
Number of mutations per branch on all branches of the SARS-CoV-2 tree with at least one mutation. The analysis used here only considers mutations on branches with four or fewer mutations and so excludes the small fraction of highly mutated branches.

